# Size matters: optimal mask diameter and box size for single-particle cryogenic electron microscopy

**DOI:** 10.1101/2020.08.23.263707

**Authors:** Toshio Moriya, Naruhiko Adachi, Masato Kawasaki, Yusuke Yamada, Akira Shinoda, Kotaro Koiwai, Fumiaki Yumoto, Toshiya Senda

## Abstract

Recently it has been demonstrated that single-particle cryogenic electron microscopy (cryo-EM) at 200 keV is capable of determining protein structures, including those smaller than 100 kDa, at sub-3.0 Å resolutions, without using significant defocus or a phase plate. However, the majority of near-atomic resolution cryo-EM structures has been determined using 300 keV. Consequently, many typical parameter settings for the cryo-EM computational image processing steps, especially those associated with the contrast transfer function, are based on the accumulated experience of 300 kV cryo-EM. We have therefore revised these parameters, established theoretical bases for criteria to find an optimal mask diameter and box size for a given dataset irrespective of acceleration voltage or protein size, and proposed a protocol. Considering the defocus distributions of the datasets, merely optimizing the mask diameters and box sizes yielded meaningful resolution improvements for the reconstruction of < 200 kDa proteins using 200 kV cryo-EM.

## Introduction

Cryogenic electron microscopy (cryo-EM) single-particle analysis (SPA) has recently emerged as a popular choice for the 3D structure determination of proteins, facilitating a deeper understanding of protein function and providing valuable information for developing medicines. The number of reported cryo-EM SPA structures at < 4 Å resolutions, including sub-2 Å, is increasing exponentially^1,2^. In an influential early SPA study, a molecular weight of 38 kDa was a predicted theoretical limit for SPA specimen size^3^, and specimen size still remains a limiting factor in practice^4–6^. Nearly 98% of these high-resolution structures have been determined using 300 kV transmission electron microscopes, but this has primarily been successful in the visualization of large protein complexes^7^. Just ∼1% of all cryo-EM reconstructions resolved to better than 4 Å resolution in the Electron Microscopy Data Bank (EMDB) are macromolecules < 200 kDa^5^.

Recently it has been demonstrated that SPA using 200 kV cryo-EM, equipped with a direct electron detector (DED), is capable of reconstructing structures of proteins smaller than 200 kDa at sub-3.0 Å^4,7,8^. The authors resolved ∼150 kDa rabbit muscle aldolase to 2.6 Å and later improved the reconstruction to 2.13 Å using conventional defocus-based SPA methods. Furthermore, biological specimens massing < 100 kDa were resolved to better than 3 Å resolution (2.72 Å for ∼82 kDa horse liver alcohol dehydrogenase and 2.8 Å for ∼64 kDa human methemoglobin). This indicates that 200 keV can be appropriate for high-resolution reconstructions of < 200 kDa proteins.

Our previous study^9^ built on the potential of 200 kV cryo-EM SPA for the high-resolution structure determination of < 200 kDa proteins. In that study, parameter settings for the image processing steps were revised, since many of the typical settings were based on the accumulated experience of large proteins (> 200 kDa) with 300 kV cryo-EMs. Such parameters include the suggested mask diameter and box size. In previous studies, deviations from the optimal values of these parameters have not prevented resolution improvement up to the current achievable levels for cryo-EM, so there have not been strong attempts to conduct methodological studies for parameter optimization. However, during the 200 kV cryo-EM SPA of 110 kDa nitrite reductase in our previous study, we noticed that the mask diameter (110 Å) and box size (256 pixels) chosen by a widely-used conventional protocol based on empirical criteria were too small. The conventional settings yielded a 0.39 Å lower resolution than the 2.85 Å final reconstruction with revised values (164 Å mask diameter and 486-pixel box with a pixel size of 0.88 Å/pixel).

This result indicated that the empirical criteria for choosing the mask diameter and box size might work properly only for 300 kV datasets of large proteins. The widely-used protocol based on the empirical criteria are, (1) set “mask diameter” slightly larger (e.g. ∼10%) than the measured diameter of the circle enclosing the largest particle view, and then (2) set “box size” at least 1.5x to 2.0x of the mask diameter, or even larger when atypically large defocus (e.g. -3.0 µm) is used. With this protocol, a smaller particle always requires a smaller box size, since the protocol adjusts the mask diameter and box size by referring only to the particle size and adjustment factors such as “slightly larger (e.g. ∼10%)” and “1.5x to 2.0x”, whose relations to the defocus are not well-defined. These adjustment factors were decided empirically and obscure the involvement of the point spread function (PSF), which is the inverse Fourier transformation of contrast transfer function (CTF).

In the current study we attempt to establish theoretical criteria for optimal mask diameter and box size for a given dataset by focusing on PSF and CTF, while keeping the computational resource requirements to a minimum for practical reasons. A too-small mask diameter or box size causes information loss in real space, and the information spread outside of the particle edge by PSF in real space must also be included for the CTF correction. According to the Nyquist–Shannon sampling theorem, a too-small box size also causes misrepresentation of CTF due to an insufficient number of sampling points even in reciprocal space. To evaluate the validity of our proposed method, the 200 kV cryo-EM datasets of two small proteins (∼64 and ∼150 kDa), available from the Electron Microscopy Public Image Archive (EMPIAR)^10^, as well as the nitrite reductase dataset (∼110 kDa) were used, in alignment with our research focus. The results of all datasets demonstrated that merely optimizing the box sizes and mask diameters yielded meaningful improvements in the ∼3.0 Å resolution range. Finally, based on our findings we proposed a protocol for determining optimal mask diameter and box size for a given dataset, irrespective of acceleration voltage or protein size, while explicitly considering the particle size, the pixel size, and the CTF parameters, including the defocus distribution.

## Results

### Theoretical analysis for mask diameter and box size

The commonly-used protocol decides the mask diameter and box size basically from the measured particle size but, as can be inferred from step (2) of the above-described protocol, better decisions could be made by considering PSF and CTF (Fig. 1). PSF is related to the mask diameter in real space. CTF is associated with the box size, which decides the number of sampling points both in real and reciprocal space, without changing pixel size and thus the Nyquist frequency. Since the adjustable ranges of other related factors are extremely limited during SPA image processing, mask diameter and box size are the most appropriate targets for optimization.

**Fig. 1.**
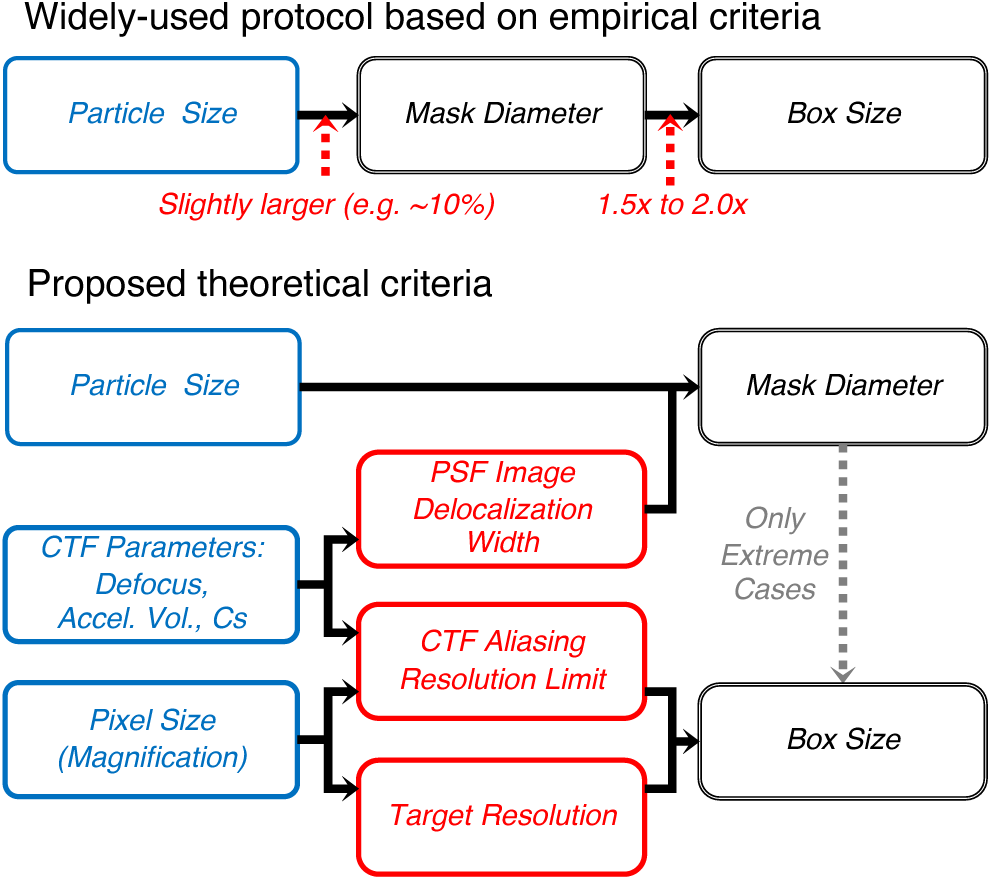
Dependencies of related parameters and criteria. Dependency charts of the parameters and criteria related to mask diameter and box size in a widely-used protocol based on empirical criteria (above), and proposed theoretical criteria (below). The base decision factors (blue) are the parameters fixed at cryo-EM imaging: the particle size, the CTF parameters (mainly, defocus, acceleration voltage (Accel. Vol.), and spherical aberration (Cs)), and the pixel size (equivalent to magnification). In the SPA image processing stage, either they are not adjustable at all or the adjustable ranges are extremely limited. The base decision factors heavily influence the theoretical criteria (red): the PSF image delocalization width, the CTF aliasing resolution limit, and target resolution (must be lower than the theoretically-achievable maximum resolution). Mask diameter and box size should be optimized to fulfill the requirements of theoretical criteria.

In cryo-EM, the information in a particle view extends beyond the “true” boundary due to the convolution effect of PSF, called “PSF image delocalization width” (hereafter, “PSF width”)^11^ (Fig. 2). This undesirable effect means that mask diameter must include the information delocalized within the PSF width for the CTF correction, in order for the CTF deconvolution to restore as much of the original information as possible. The CTF correction is embedded in many SPA processing steps and is a key to achieving near-atomic resolution, especially in 3D refinement. A larger defocus value makes the PSF width wider and so requires a larger mask diameter. That is, the CTF parameters influence the PSF width, and in turn the PSF width and the particle size together determine the required mask diameter. Here, the decision of optimal mask diameter is dominated by the particle size, but it has to be adjusted depending on the defocus value to compensate for the PSF width.

**Fig. 2.**
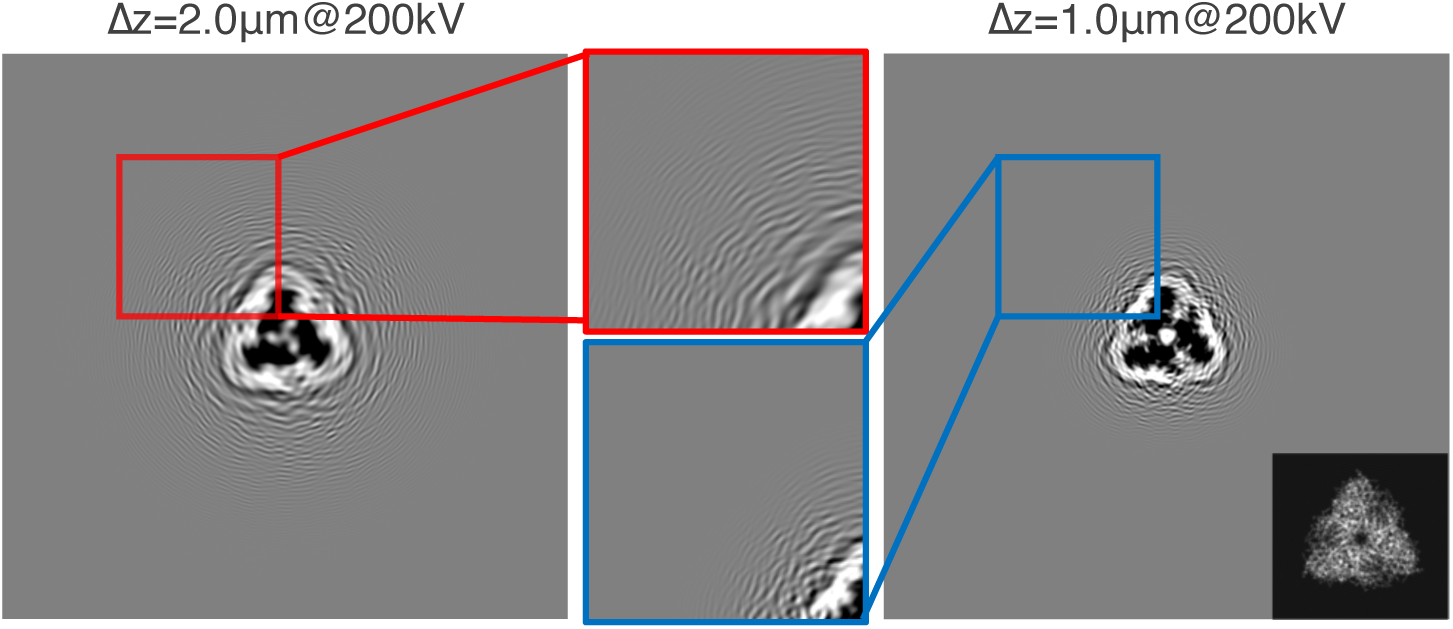
PSF image delocalization width of particle view. The information extension of particle views due to PSF in real space. The CTFs were simulated at two different defocus values: 2.0 µm (red) and 1.0 µm (blue) using an acceleration voltage of 200 kV, spherical aberration 2.7 mm, B-factor 80.0 Å^2^, pixel size 1.0 Å/pixel, and box size 512 pixels. The simulated particle images of nitrite reductase in real space after applying the CTF model are shown on the left and right, with the original 512-pixel box size and zoomed-in views in the middle. The simulated image, before applying the CTF model, is shown at the right bottom. The ripple patterns are the information extended beyond the original boundary of the particle view.

The problems associated with the box size are rather complex and occur in reciprocal space. The main problem is aliasing artifact that can emerge in reciprocal space because a discrete Fourier transformation with a too-small box size in real space (meaning an small number of sampling points in real space) will yield an insufficient number of sampling points in reciprocal space (Fig. 3). Note that the number of sampling points in reciprocal space must be the same as the number in real space, in the discrete Fourier transformation. Therefore, when applying the Nyquist–Shannon sampling theorem (hereafter, “sampling theorem”) to reciprocal space, too-rapid oscillation part in CTF, whose frequency exceeds the Nyquist frequency (i.e., the inverse of twice the reciprocal pixel size), creates CTF aliasing artifacts. A larger defocus value makes the oscillation more rapid in the high-frequency range (Fig. S1). Furthermore, the oscillations of 200 kV CTF curves are more rapid than those of 300 kV curves when the defocus values are the same, due to the different electron wavelengths (Fig S1). Therefore, the box size must be large enough to represent the CTF oscillation in reciprocal space correctly. Otherwise, the CTF aliasing artifacts interfere with the CTF correction performed on the discrete Fourier transform of each particle image in reciprocal space.

**Fig. 3.**
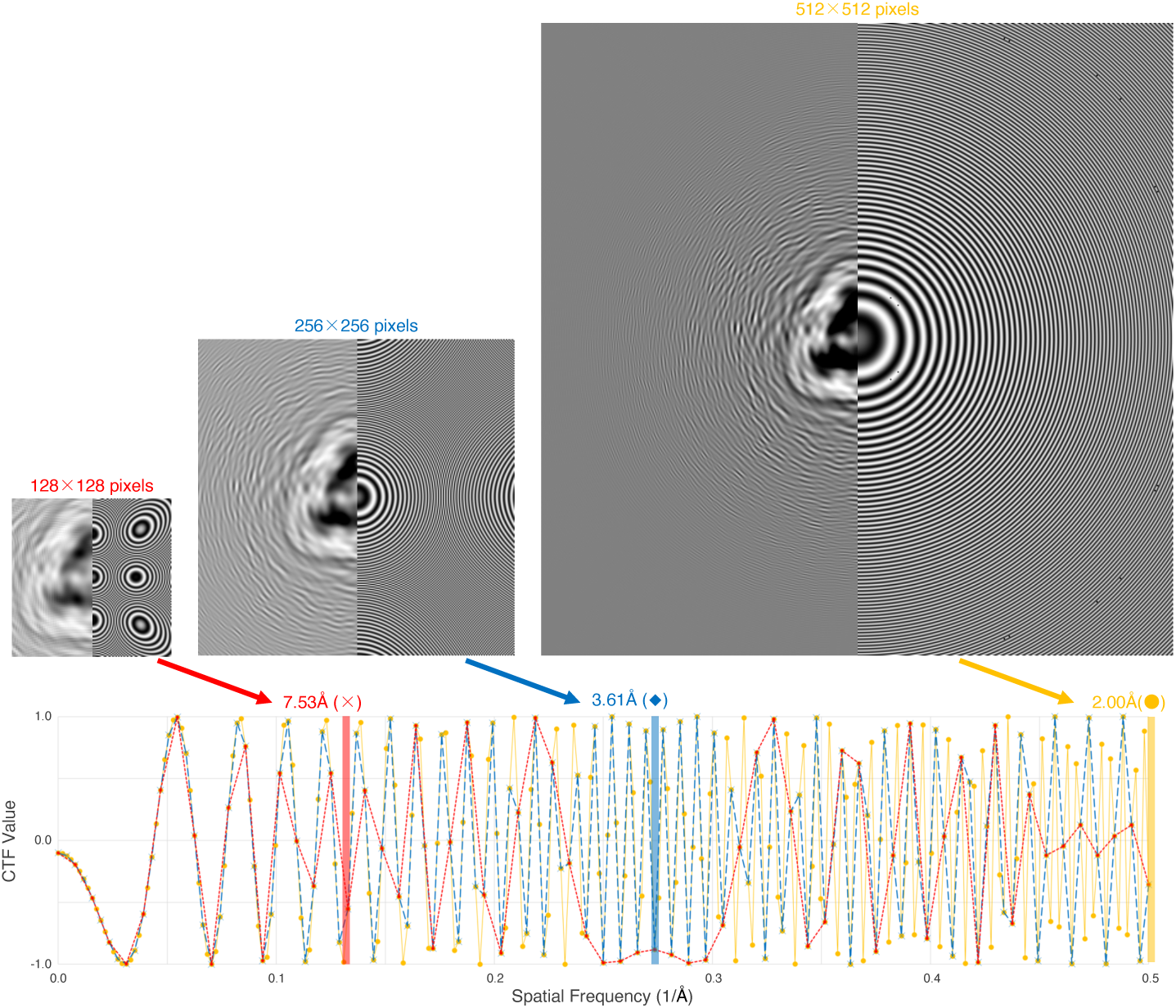
CTF aliasing frequency limit and box size. The relationship between the CTF limit and box size. The CTF curves were simulated for the box sizes of 512, 256, and 128 pixels, acceleration voltage 200 kV, spherical aberration 2.7 mm, defocus 2.00 µm, and pixel size 1.0 Å/pixels. (Top) In images derived from each box size, right is the 2D CTF model and left is the simulated particle image of nitrite reductase in real space after applying the CTF model. The CTF aliasing artifacts, which are patterns other than the concentric circles of Thon rings, are clear in the 256- and 128-pixel box sizes. (Bottom) 1D curves of the corresponding CTF models of all the box sizes. The vertical lines indicate the CTF limits: 7.53 Å for the 128-pixel box size (red), 3.61 Å for 256-pixel (blue), and 2.00 Å (Nyquist) for 512-pixel (yellow). Beyond the CTF limits, the CTF curves of the 128-and 256-pixel box sizes diverge from the aliasing-free CTF curve of the 512-pixel box size.

Penczek *et al*. pointed out this problem and defined the “CTF aliasing resolution limit” (hereafter, “CTF limit”)as the highest frequency up to which a CTF model can be represented correctly with a given box size^12^. A program that computes the CTF limit is available as the *ctflimit* function in the morphology.py Python module of the SPARX/SPHIRE software package^13,14^. It ensures that no reciprocal-space aliasing in a CTF model occurs at lower than its output frequency value for the inputs of box size, CTF parameters (i.e., defocus value, spherical aberration (Cs), and acceleration voltage), and pixel size. The pixel size (defined in real space and equivalent to magnification) also determines a theoretically-achievable maximum resolution (i.e., the inverse of the Nyquist frequency or twice the pixel size). The user-defined target resolution (hereafter, “target resolution”), which the user expects to achieve for the final result, must be lower than the theoretical limit. Therefore, the box size must be large enough so that the CTF limit is higher than the target resolution (hereafter, “CTF-limit box size”). Importantly, the CTF-limit box size would have no relation to the particle size (Fig. 1). When large defocus values are used, a larger box size is required even for a small protein, meaning that a smaller box size is not necessarily optimal for a smaller particle.

To address the issues inherent in basing a protocol on empirical criteria, we conducted mask diameter and box size experiments. Strictly speaking, the box size must also be larger than the mask diameter so that the box size does not cut out any information necessary for the CTF correction, which must be enclosed by the mask diameter (Fig. 1). It is exceptional for the mask dimeter to be larger than the CTF-limit box size, but such cases do happen with exceptionally large proteins, when the defocus range is limited to atypically small values, or with an extremely low target resolution. In typical cases, the optimal box size is the CTF-limit box size, so this dependence was not considered explicitly in the current study.

### Mask diameter experiment

The mask diameter experiment was conducted first. Our previous study^9^ demonstrated that the optimal box size was the CTF-limit box size, based on the maximum defocus value in the dataset (2.64 Å; 3/2 Nyquist resolution for 0.88 Å/pixel pixel size), and was far larger than the particle size (∼4.3 times) as well as larger than the optimal mask diameter (∼2.6 times). Therefore, the CTF-limit box size for the maximum defocus was a safe choice to ensure both that it would be larger than the yet-unknown optimal mask diameter and that all particle images would be free from CTF aliasing artifacts. By basing the box size on the CTF limit of the maximum defocus in this experiment, the independent effect of mask diameter variation could be obtained, and the interpretation of the results would be simplified.

The criteria to determine PSF width are tricky, since there are no obvious ways to calculate the width of the image delocalization even for a single defocus value. Therefore, we gave up on an analytical solution and decided to employ a numerical solution, which utilized an intermediate result of the SPA steps. We have been observing that the 3D density map of 3D refinement usually has negative densities surrounding positive particle density volume at the center (Fig. 4 left panels). Since all popular SPA algorithms use the zero-background normalization (set the background vitrified ice density to zero), we hypothesized that the negative density volume is a part of the object and the extent of the negative density is in fact the total extent of the PSF influence, due to the imperfect CTF correction.

**Fig. 4.**
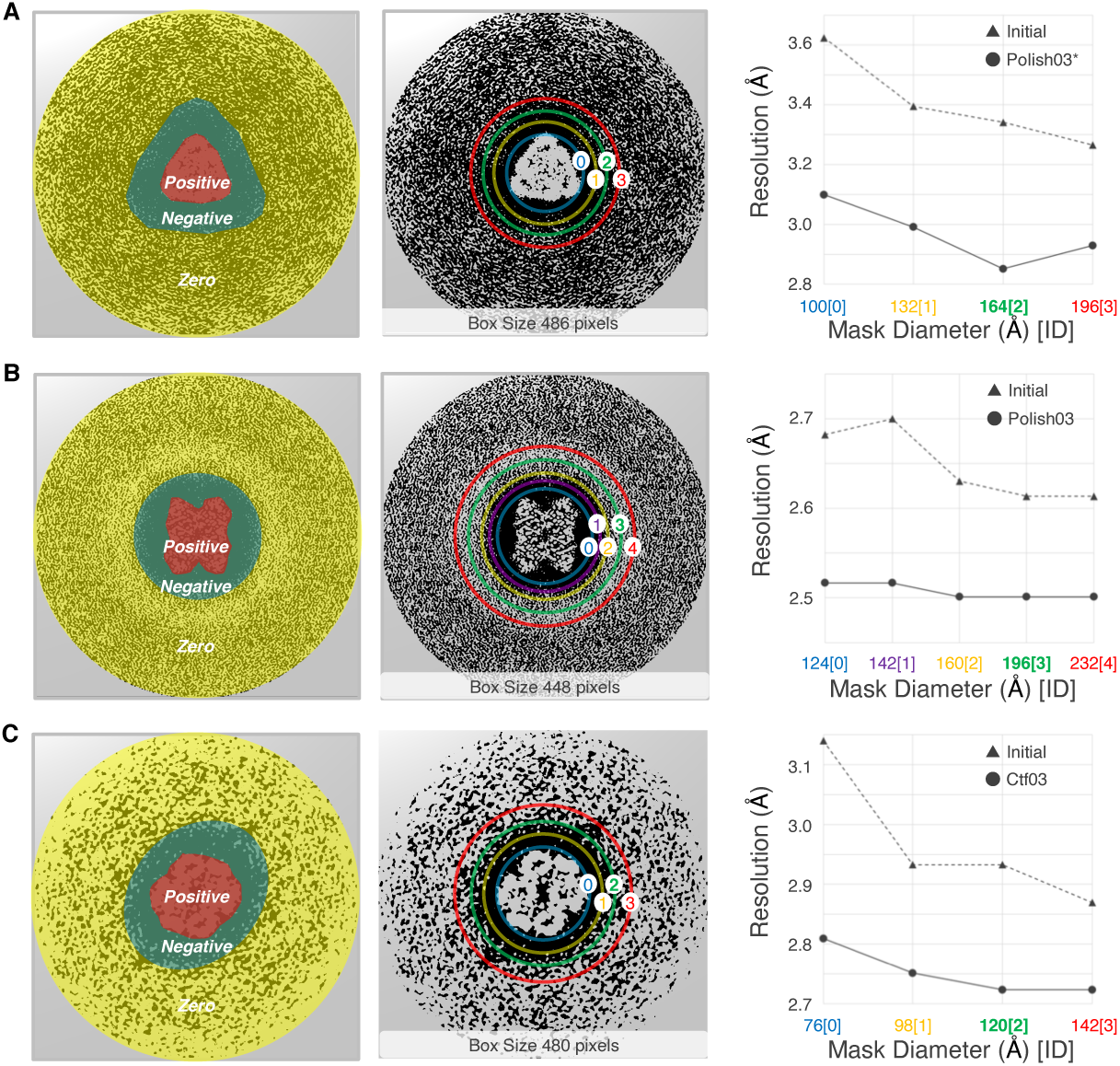
Results of the mask diameter experiment. The results of the mask diameter experiment with (A) nitrite reductase, (B) muscle aldolase, and (C) methemoglobin. Left panels depict the negative density areas in the central sections, orthogonal to the z-axis direction, of the 3D cryo-EM maps obtained by the 3D refinement steps at the end of the particle screening processes. In each map, the outermost is the zero density area (yellow); the negative density area (green) surrounds the positive density area (red). Middle panels indicate the mask diameters used in each dataset. Right panels are the plots of the resolutions relative to the mask diameter variations. The dotted line is the experiment at the initial 3D refinement step (“Initial”); solid is the experiment at the best 3D refinement step obtained during the three cycles of CTF refinement and Bayesian polishing. The *x*-axis is the mask diameter in Å and mask dimeter ID corresponding to the number in the middle panels. The *y*-axis is the 0.143 FSC resolution in Å.

To test the hypothesis of whether the extent of the negative density area diameter is really the optimal mask diameter, multiple 3D refinements of Relion3^15^ were executed using different values for the mask diameter option while keeping the other input parameter settings the same. The 200 kV cryo-EM datasets of ∼110 kDa native nitrite reductase (EMD-0731)^9^, ∼160kDa rabbit muscle aldolase (EMPIAR-10181)^7^, and ∼64 kDa human methemoglobin (EMPIAR-10250)^4^ datasets were used (Table 1). Prior to the experiment, each dataset was cleaned using multiple executions of the Relion3 2D/3D classification, closely following the original publication. In this way we obtained a stack of particle images with the highest resolution information (see also “Image processing for 3D reconstruction” subsection in Methods). For each cleaned dataset, the 3D refinement was then repeated using four or five different mask diameters while keeping the box size constant, as described above (Table 2). Additionally, the effect of the Relion3 CTF refinement and Bayesian polishing on the optimal value of mask diameter was examined. Using the optimal mask diameter determined for each dataset, three cycles of the CTF refinement and Bayesian polishing were executed to improve the CTF estimations by refining per-particle defocus, correcting beam tilt, and compensating for beam-induced motion of each particle image. Then, the same procedure of the mask diameter variation was repeated with the best 3D refinement step obtained during the cycles. To compare the processing times of different mask diameters, all calculations for the methemoglobin dataset were executed with the same single desktop computer equipped with four graphics processing unit (GPU) cards.

**Table 1.**
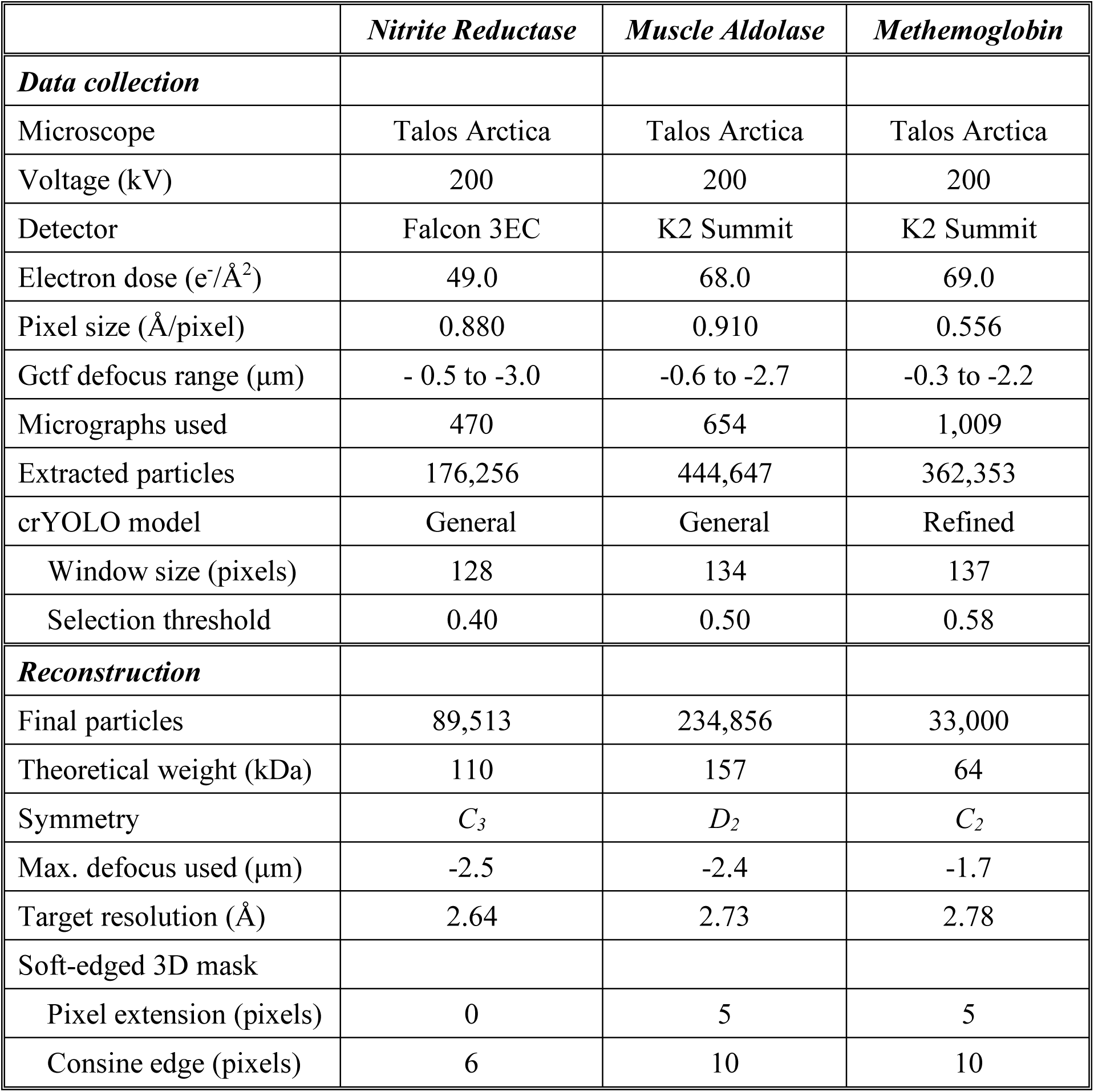
Cryo-EM data collection and reconstruction

**Table 2.** Parameter settings of the mask diameter (mask Ø) experiment

|  | *Mask Ø (Å)* | *Mask Ø / Positive Ø* | *Note* |
| --- | --- | --- | --- |
| ***Nitrite Reductase*** | ***(0) 100*** | 1.00 | Postive Ø |
| *Pixel size 0.88 Å/pixel* | ***(1) 132*** | 1.32 |  |
| *Box size 486 pixels (429 Å)* | ***(2) 164*** | 1.64 | Negative Ø |
|  | ***(3) 196*** | 1.96 |  |
| ***Muscle Aldolase*** | ***(0) 124*** | 1.00 | Postive Ø |
| *Pixel size 0.91 Å/pixel* | ***(1) 142*** | 1.15 |  |
| *Box size 448 pixels (408 Å)* | ***(2) 160*** | 1.29 |  |
|  | ***(3) 196*** | 1.58 | Negative Ø |
|  | ***(4) 232*** | 1.87 |  |
| ***Methemoglobin*** | ***(0) 76*** | 1.00 | Postive Ø |
| *Pixel size 0.556 Å/pixel* | ***(1) 98*** | 1.29 |  |
| *Box size 480 pixels (267 Å)* | ***(2) 120*** | 1.58 | Negative Ø |
|  | ***(3) 142*** | 1.87 |  |

Fig. 4 summarizes the results (see also Tables S1-S3). All datasets showed a similar trend both before and after the CTF Refinement and Bayesian Polishing cycles. The smallest mask diameters, which were very close to the boundaries of the positive and negative density areas, always resulted in the lowest resolutions. The highest resolutions were obtained with diameters around the same size or larger than the negative density areas. Using diameters larger than the negative density area did not appear to improve resolution further, indicating that improvement saturates beyond the negative density size. Therefore, the diameter closest to the negative-zero density boundary was optimal in terms of the resolution. With all datasets, differences between the best and worst resolutions were much larger at the initial 3D refinement step than the best 3D refinement during the CTF Refinement and Bayesian Polishing cycles (0.36 Å vs. 0.25 Å for nitrite reductase, 0.09 Å vs. 0.02 Å for aldolase and 0.27 vs. 0.09 Å for methemoglobin). The processing times of the 3D refinements with all mask diameters were similar both before and after the CTF Refinement and Bayesian Polishing cycles, showing no obvious trend relative to the mask diameter variation (Table S4). These results were consistent with our hypothesis that the negative density area mainly reflects the total extent of the PSF influence due to imperfect CTF correction, and that the inclusion of this area in the mask diameter indeed improves resolution in 3D reconstructions.

### Box size experiment

For computation of the CTF-limit box size, a difficulty of establishing criteria based on the CTF limit is that it requires a single defocus value. However, a cryo-EM dataset has to be taken with a wide range of defocus values (Fig. S2) to compensate for the zero-crossings of CTFs where no information is transferred (Fig. S1). It is evident that the safest method is to calculate the box size where the CTF limit of the “maximum defocus value” in the dataset is higher than the target resolution. This ensures that all particle images are free from CTF aliasing artifacts for the target resolution. However, this criterion can introduce practical difficulties related to computation hardware limitations. A larger defocus value requires a larger box size, but calculation with an excessively large box size might be impossible because of insufficient memory size or impractically long calculation time. Therefore, in practice it is desirable to use a smaller defocus value for the calculation of the CTF limit. To this end, the present study aimed to find a defocus value which is smaller than the maximum but still represents the “whole” dataset and produces the same level of resolution.

We conducted the box size experiment (see Methods) with Relion3^15^ using the same cleaned datasets as in the mask diameter experiment. Three defocus values were tested: the 100^th^ (maximum), ∼75^th^, and ∼50^th^ percentile value of the defocus distribution in each dataset (Table 3 and Fig. S2). For the choice of the target resolutions, the 3/2 Nyquist resolution (three times the pixel sizes) was preferred in the current study because this resolution limit criterion is relatively conservative but still practically plausible^16,17^. When the pixel size is small (e.g., 0.5 Å/pixel), the 3/2 Nyquist resolution can be too high to keep the CTF-limit box size within a practical level. In this case, the target resolution was selected based on the resolutions achieved by the original studies. For each box size of each dataset, the 3D refinement was followed by three cycles of the CTF refinement and Bayesian polishing in Relion3. To measure the resolution improvement, the 3D refinement was performed after each processing step. The padding option was turned off (i.e., set “Skip padding” to “Yes”) for all of the 3D refinement runs. For all the steps in this experiment, the particle mask diameter (“*Mask Ø”*) was kept constant to the optimal value obtained in the mask diameter experiment, to ensure that the effect of the box size variation would be independent from the information loss due to the PSF width. To compare the processing times relative to different box sizes, all calculations of the methemoglobin dataset were again executed with the same desktop computer, as in the mask diameter experiment.

**Table 3.**
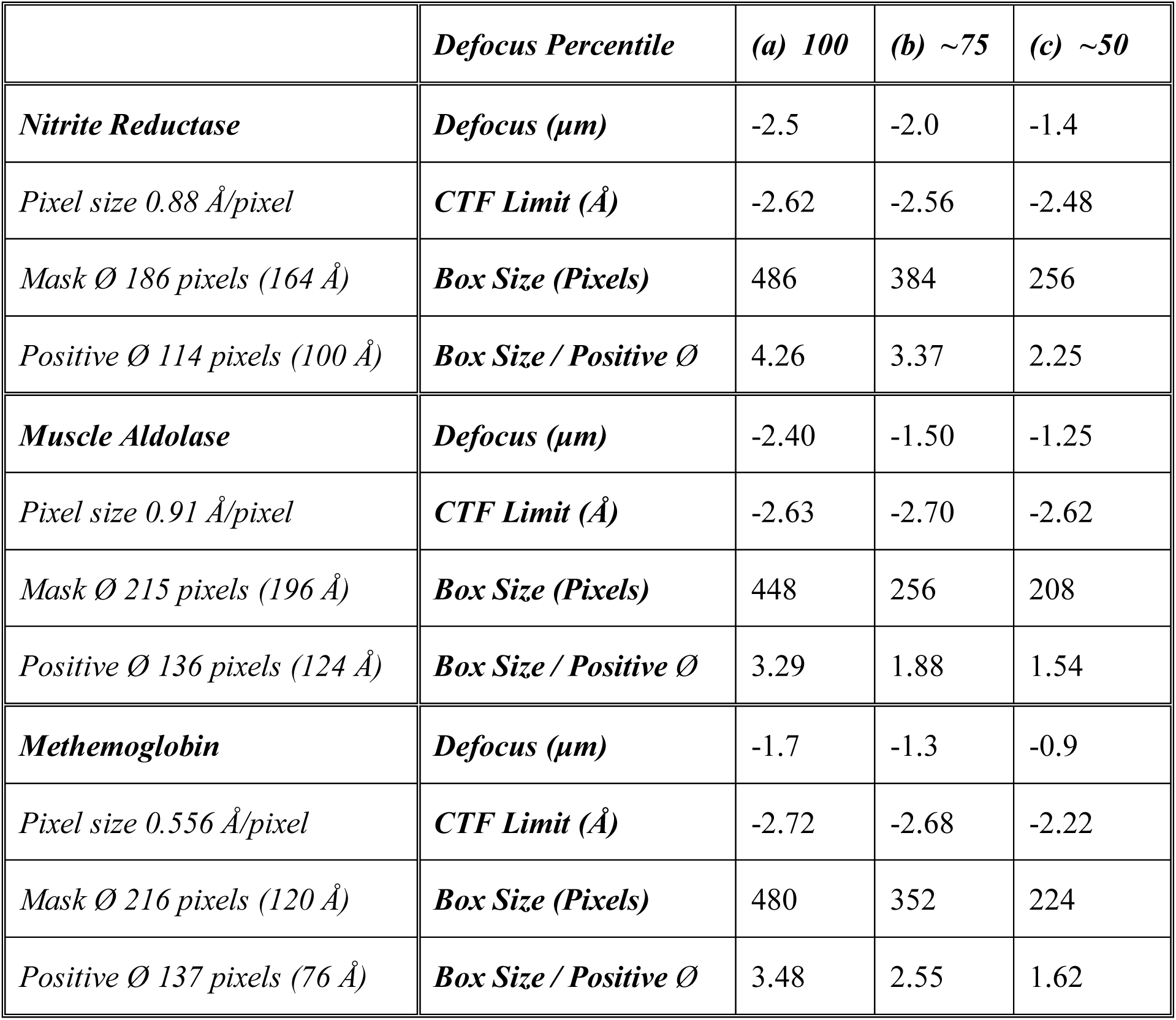
Parameter settings of the box size experiment

The results of all datasets are summarized in Fig. 5 (see also Tables S5-S7). With native nitrite reductase, the resolutions of all the box size settings were almost identical at the initial 3D refinement step. During the cycles of the CTF Refinement and Bayesian Polishing, resolutions improved with trends similar to each other regardless of box size. However, the improvement of the 486- and 384-pixel box sizes, decided by the CTF limits (2.62 and 2.56 Å) at the 100^th^ and ∼75^th^ percentile defocus values (−2.5 and -2.0 µm), was much larger than that of the 256-pixel box size (CTF limits 2.48 Å with the ∼50^th^ percentile defocus values of -1.4 µm). The 486-pixel box showed slightly larger improvement than the 384-pixel box size. This suggests that the optimal box size would be based on a CTF-limit of defocus value somewhere between the 75^th^ and 100^th^ percentile of the distribution for the nitrite reductase dataset.

**Fig. 5.**
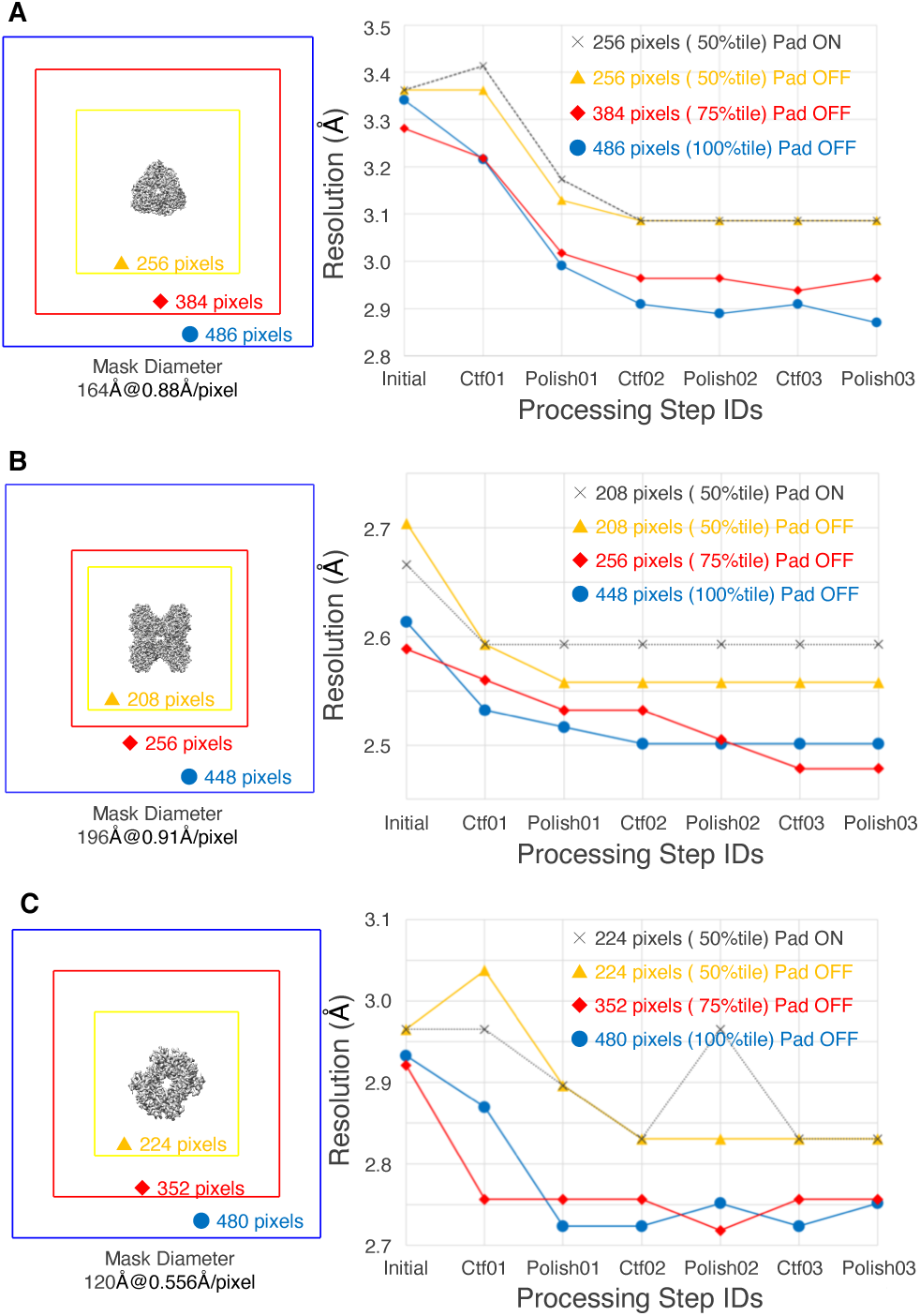
Results of the box size experiment. The results of box size experiments with (A) nitrite reductase, (B) muscle aldolase, and (C) methemoglobin. Left panels depict box sizes (in pixels) used in each dataset relative to the positive density volume of the particle. Right panels are the plots of the resolution improvement during three cycles of CTF refinement and Bayesian polishing, starting from the initial 3D reconstructions. The *x*-axis is the step ID: “Initial” is initial 3D refinement, “Ctf01” is 1^st^ CTF refinement, “Polish01” is 1^st^ Bayesian Polishing, “Ctf02” is 2^nd^ CTF refinement, “Polish02” is 2^nd^ Bayesian Polishing, “Ctf03” is 3^rd^ CTF refinement, and “Polish03” is 3^rd^ Bayesian Polishing. The *y*-axis is the 0.143 FSC resolution in Å.

The box size experiments with the EMPIAR datasets produced similar results; the resolutions improved with almost the same trend regardless of the box size during the cycles of the CTF Refinement and Bayesian Polishing. In general, the resolution difference due to the optimization of the box size was larger after the CTF Refinement and Bayesian Polishing cycle than in the initial 3D refinement, except the muscle aldolase dataset. With this dataset, the resolution of the initial 3D refinement with a 208-pixel box size (∼50^th^ percentile defocus CTF limit) was much lower than the 448- and 256-pixel box size settings (0.12 Å difference). Comparing box sizes based on the ∼75^th^ and 100^th^ percentile defocus CTF limits, the improvement trends and resolution values were almost identical to each other in both EMPIAR datasets. Also, processing times with the methemoglobin dataset were as expected; the larger the box size, the longer the processing time (Table S8). This indicates that the ∼75^th^ percentile defocus CTF limit can give us a box size very close to optimal.

Additionally, we executed the CTF Refinement and Bayesian Polishing cycles of the smallest box size setting with each dataset, using the same input parameter settings, except that the padding option of the 3D Refinement was turned on. With this option, the sizes of particle images were internally increased by padding with zeros (2 times the original box size). The resulting resolution improvement curves were almost identical with the padding option ON and OFF (Fig. 5 left panels, gray and yellow curves), although the processing with the option ON took much longer (∼1.44 times) (Table S8). This indicates that the padding option does not compensate for a too-small box size when the CTF limit is considered.

### Direct comparisons between empirical and proposed theoretical criteria

In the experiments of mask diameter and box size, the best resolution of the nitrite reductase reconstruction was 2.85 Å (Fig. S3). For the EMPIAR datasets, we achieved the resolutions slightly better than or comparable to those of the original publications^4,7^; 2.48 Å for muscle aldolase (Fig. S4) and 2.72 Å for methemoglobin (Fig. S5).

To check whether these best results were indeed better than those obtained with the widely-used empirical criteria, the box size experiment was performed again using the mask diameter and box size determined by empirical criteria. Direct comparisons between empirical criteria and proposed theoretical criteria for the selection of mask diameters and box sizes indeed showed the superiority of our proposed criteria for all datasets (Fig. 6). Interestingly, the resolution change of the methemoglobin dataset was far more unstable when using the empirical mask diameter and box size compared with our optimal values (Fig. 6C). The CTF refinement steps yielded prominently worse resolutions than the Bayesian polishing steps with the empirical values. This might be another benefit of using our proposed theoretical criteria, as it allows a smooth convergence of the resolution improvement during the CTF refinement and Bayesian polishing cycles.

**Fig. 6.**
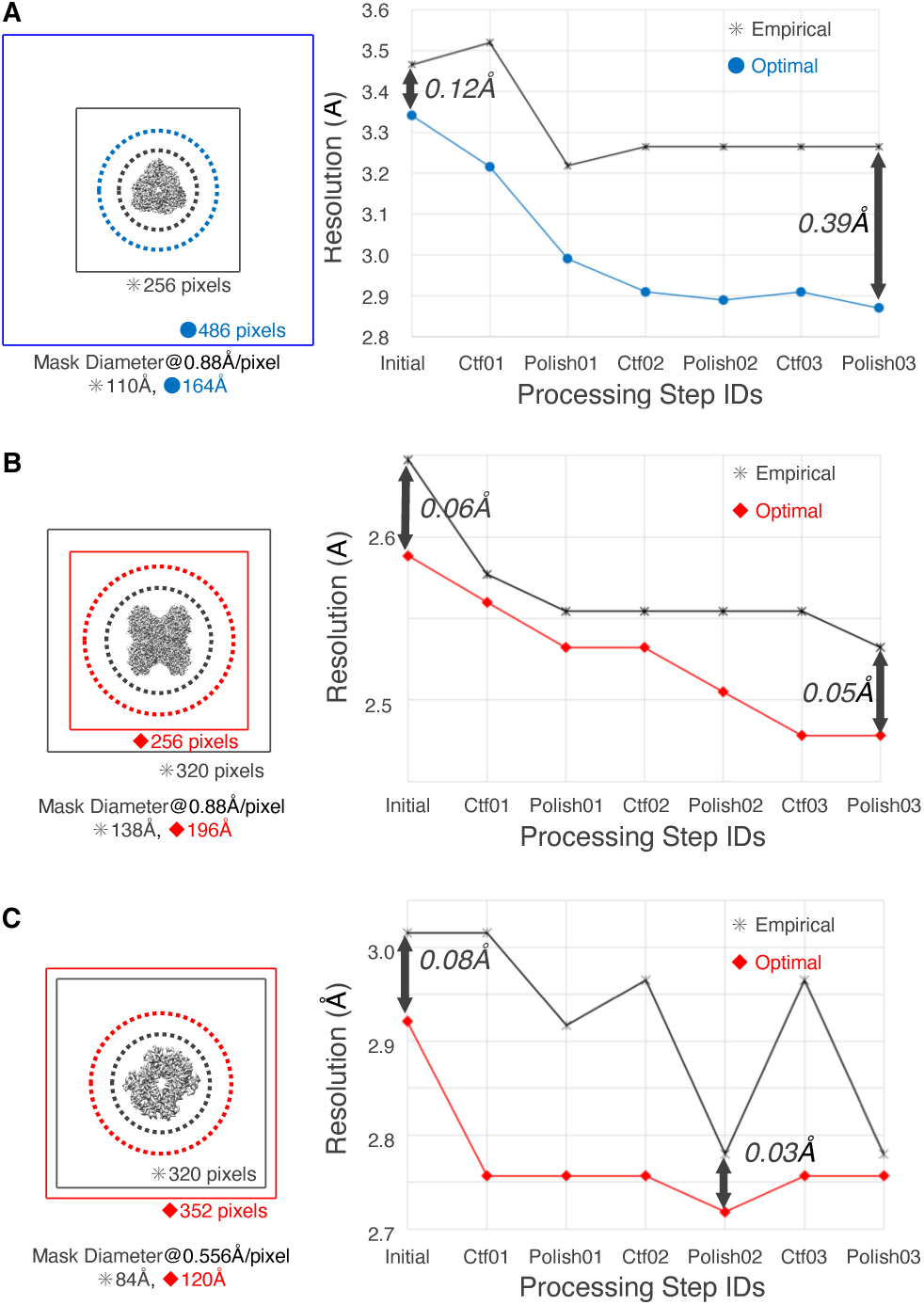
Comparisons between empirical and optimal settings. The results of direct comparisons between empirical criteria (“Empirical”) and proposed theoretical criteria (“Optimal”) for the selection of mask diameter and box size. (A) nitrite reductase, (B) muscle aldolase, and (C) methemoglobin. The layout, labels, colors, and graph axes of the panels are the same as Fig. 5. The double-headed arrows indicate the resolution differences between the two criteria at the initial 3D refinement, and at the best 3D refinement step obtained during the three cycles of CTF refinement and Bayesian polishing.

### Proposal of optimization protocol

Based on our analysis, we propose the following protocol to find optimal mask diameter and box size (Fig. 7). (1) Decide the target resolution *f*_*target*_ for final 3D reconstruction (the resolution one wants to achieve). (2) Find the *∼*75^th^ percentile value *Δz*_*75*_ of the defocus distribution of the dataset. (3) Compute the CTF-limit box size *B*_*f*_ based on CTF limit *f*_*limit*_ for *Δz*_*75*_ where *f*_*limit*_ ≤ *f*_*target*_. (4) Measure the diameter of the positive particle density area of the longest particle view *Φ*_*+*_. (5) Calculate initial mask diameter *P*_*init*_ at twice the measured particle diameter (i.e., *2Φ*_+_). (6) Calculate the box size *B*_*p*_ so that *P*_*init*_ is 95% of this box size (i.e., *P*_*init*_ / *0*.*95*), to make sure that the particle image has some margin for zero-background volume, so that the boundary of the negative particle density area will be easily found in step 9. (7) Choose the larger of *B*_*f*_ and *B*_*p*_ as the optimal box size *B*_*opt*_. (8) Execute initial 3D refinement using *B*_*opt*_ for the box size and *P*_*init*_ for the mask diameter, to get the 3D density map *V*_*init*_. (9) In *V*_*init*_, measure the diameter of the negative particle density volume *Φ-*surrounding the positive particle density volume. (10) Choose *Φ-*as the optimal mask diameter *Φ*_*opt*_ for the subsequent 3D refinement and 3D classification steps.

**Fig. 7.**
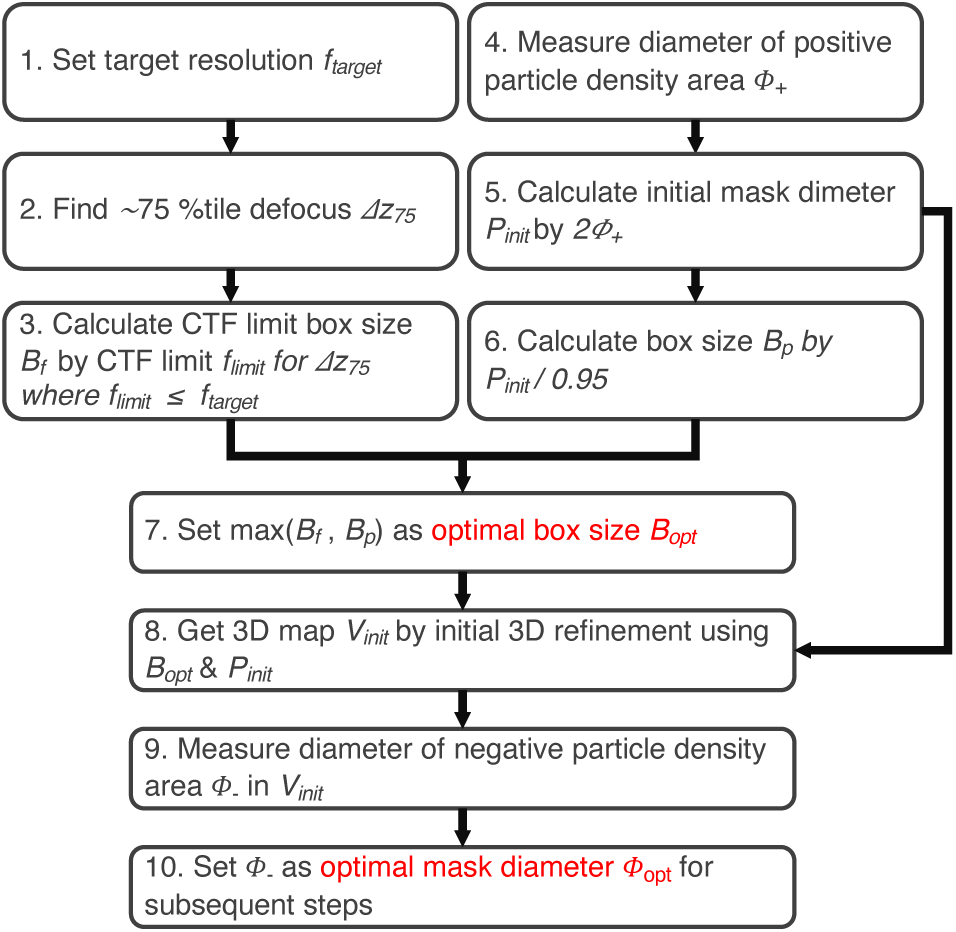
Flowchart of the proposed protocol. The flowchart of the proposed protocol for finding optimal mask diameter and box size. The protocol was established based on the findings from the mask diameter and box size experiments.

## Discussion

The current study demonstrates the importance of choosing appropriate values for both the mask diameter and box size, especially in the case of 200 kV cryo-EM SPA of a small protein (< 200 kDa). In real space, the information spread outside of the particle edge by PSF^11^ also needs to be included within the mask diameter, meaning that the optimal mask diameter is dependent on not only the particle size but also the defocus range of the dataset. For a small protein, the choice of box size is likely not dominated by the particle size but rather by the CTF aliasing frequency limit^12^. Thus, even for a small protein, one still has to consider a larger mask diameter and box size than the empirical criteria would suggest, as the empirical criteria are based on 300 kV cryo-EM studies of large proteins. Although 200 kV cryo-EM SPA for small proteins was examined here, the proposed optimization protocol is applicable to any acceleration voltage and to any size of protein.

The most significant finding of this study is that the size of the negative density volume surrounding the positive density volume of the target particle gives a very good measurement of the optimal mask diameter. Our results demonstrate that the negative density volume has to be considered as a part of the foreground object (i.e., particle) instead of the background (i.e., pure vitrified ice), due to the PSF effect and imperfect CTF correction. Interestingly, the optimal mask diameters of all datasets were within the range of 1.5 to 2.0 times the measured diameter of the positive particle density (“Mask Ø / Positive Ø” in Table 2). This suggests that the conventional settings may be more suitable for the mask diameter than for the box size. Therefore, the conventional settings can be used as an initial approximation of optimal mask diameter until the initial 3D refinement. In conclusion, not only the box size but also the mask diameter is key to retaining the information necessary for the CTF correction, and the mask diameter should be much larger than what is used in conventional practice.

The current study also demonstrates that the optimal box size can be calculated with a CTF limit of defocus value at approximately the 75^th^ percentile. This indicates the statistical property of CTF limits; aliasing artifacts in a small number of micrographs become negligible because they can be averaged out. Although the safest choice for the optimal box size is based on the CTF limit of the maximum defocus value in a dataset, this ∼75^th^ percentile defocus criterion is still important in practice since an excessively large box size can demand too much memory and processing time. The use of small defocus values in cryo-EM imaging is also recommended, because the necessity for a large box size and mask diameter will be eliminated. However, the use of too-small defocus values (less than 0.3-0.5 µm) causes the CTF parameter estimations to become increasingly unreliable^18^. Notably, our results also indicate that increasing the particle image dimensions with the zero-padding internally in the 3D refinement cannot be used as a substitution for optimizing the box size.

After the “Resolution Revolution”^19^, near-atomic resolution has become attainable using the cryo-EM datasets. For the reconstruction of such information, the preciseness of CTF estimation is critical^18,20^. Also, many algorithms using more precise CTF models have recently yielded noticeable resolution improvement. They include per-particle CTF estimation, Ewald sphere correction, and beam tilt correction^15,21–24^. However, although a given algorithm may be capable of high-precision CTF correction, it will not work if insufficient information is input. A too-small mask diameter and/or box size essentially cuts out or deteriorates some of the information necessary for CTF correction. In this study, the per-particle defocus estimation and beam tilt correction successfully improved the resolutions in all datasets. Others have reported different results in per-particle defocus estimation and beam tilt correction trial^4^, but our results imply that mask diameter or/and box size might not have been optimal in these studies. Considering the effectiveness of more precise CTF models, including a recently developed algorithm which corrects higher-order aberrations^8,25^, the use of optimal mask diameter and box size is also expected to become increasingly crucial for these algorithms to work.

## Methods

### Datasets

To assess the validity of the proposed theoretical criteria for finding the optimal values of mask diameter and box size, three datasets of small proteins (< 200kDa) collected with 200 kV acceleration voltage were used: ∼110 kDa native nitrite reductase (EMD-0731)^9^, ∼160kDa rabbit muscle aldolase (EMPIAR-10181)^7^, and ∼64 kDa human methemoglobin (EMPIAR-10250)^4^. The detailed descriptions of sample handling, protein purification, cryo-EM grid preparation, and data acquisition can be found in previous publications. All the relevant parameters of the data collection are summarized in Table 1. All the cryo-EM datasets were collected by Talos Arctica transmission electron microscope (Thermo Fisher Scientific, Inc.) operating at 200 kV equipped with either a Falcon 3EC DED (Thermo Fisher Scientific, Inc.) or K2 summit DED (Gatan, Inc.).

### Software

Motion correction and dose-weighting were performed using the MotionCor2 frame alignment program^26^. The CTF parameters were estimated using Gctf^27^ and CTFFIND4^28^. The particles were picked using SPHIRE-crYOLO^29^. The *ctflimit* function^12^ implemented in a Python module, morphology.py, of SPARX/SPHIRE^13,14^ was used to calculate the smallest box size that ensures no CTF aliasing in the reciprocal space, up to the target resolution for a given defocus value. Relion3^15^ was used for all the other SPA steps: reference-free 2D classification, *ab initio* reconstruction, 3D classification, 3D refinement, CTF refinement for the refinement of per-particle defocus and beam tilt, and Bayesian polishing for beam-induced motion corrections^30^. After each of the 3D refinements, “gold-standard” FSC resolution with a 0.143 criterion^31^ in Relion3 was used as a global resolution estimation with phase randomization, to account for possible artifactual resolution enhancement caused by solvent mask^32,33^. The local resolutions of the 3D cryo-EM maps were estimated using RELION3’s own implementation. UCSF Chimera^34^ and *e2display*.*py* of EMAN2^35^ were used for the visualization of the output 2D/3D images.

### Image processing for 3D reconstruction

Prior to the mask diameter and box size experiments, all the datasets were cleaned with multiple runs of 2D/3D classification by closely following the original publications of nitrite reductase^9^, muscle aldolase^7^, and methemoglobin^4^. Table 1 summarizes important parameters related to the particle image screening processes of all the datasets. For each dataset, a stack of particle images was initially extracted from dose-weighted sum micrographs in either fully- or semi-automated fashion using SPHIRE-crYOLO. For fully-automated particle picking, the general model (neural network pretrained by the developer with training sets consisting of 38 real, 10 simulated, and 10 particle-free datasets on various grids with contamination) was used without any additional training (“General” for “crYOLO model” in Table 1). For semi-automated particle picking (“Refined” for “crYOLO model”), the general model was refined by fine-tuning only the last convolutional layers specifically for the dataset, while keeping the weights of all other layers fixed. After digital screening of the particle stack with multiple 2D and 3D classifications at various processing stages, the selected particles were subject to initial 3D refinement in the experiments of the mask diameter and box size. Here, the particles with large defocus were excluded to keep box size at a practical level (“Max. defocus used (µm)” in Table 1 and Fig. S2). For nitrite reductase and muscle aldolase datasets, the target resolutions (“Target resolution (Å)” in Table 1) were set to the 3/2 Nyquist resolutions. For methemoglobin, target resolution was selected based on the resolution achieved by the original study, due to the small pixel size of 0.556 Å/pixel.

Parameter settings for the mask diameter experiment are listed in Table 2. Four or five different mask diameters were used. For each dataset, the initial 3D refinement was repeated using each mask diameter while keeping the box size based on the CTF limit of the maximum defocus value constant and all other input parameters the same. The obtained resolutions of these runs were compared. To see the effect of the quality of the CTF parameters, three cycles of CTF refinement and Bayesian polishing were executed to improve the CTF estimations using the optimal mask diameter determined for each dataset. Following this, the same procedure of mask diameter variations was repeated using the best 3D refinement obtained during the CTF refinement and Bayesian polishing cycles. For the nitrite reductase dataset the 3^rd^ Bayesian polishing step was the best, so this was repeated with electron dose adjustment by removing the last 8 movie frames and using the output for the mask diameter experiment. The 3^rd^ Bayesian polishing step was also the best with the muscle aldolase dataset. For the methemoglobin dataset, the 3^rd^ CTF refinement step was the best.

Table 3 shows the parameter settings used in the box size experiment. These box sizes ensured that all the CTF limits calculated with the maximum values, ∼75^th^ percentile values, and ∼50^th^ percentile values of the defocus ranges (Fig. S2) were higher than the target resolution (“Target resolution (Å)” in Table 1). The optimal values obtained in the mask diameter experiment were used as mask diameters (“*Mask Ø”*) for all the subsequent processing steps in this experiment. For each box size, the initial 3D refinement with a soft-edged 3D mask was performed by imposing symmetry and using no padding (i.e. set “Skip padding” to “Yes”). Then, three cycles of CTF refinement and Bayesian polishing were executed to refine per-particle defocus, beam tilt, and beam-induced motion corrections. The number of repeats ensured no further improvements. To measure the degree of resolution improvement, 3D refinement (symmetry imposed but without padding) with a soft-edged 3D mask and with solvent-flattened FSCs options was used after each CTF refinement and each Bayesian polishing step. In addition, the smallest box size setting of each dataset was repeated, using the same input parameter settings, except that the padding option of the 3D refinement was turned on to internally increase the box size twice by padding with zero.

For the direct comparisons between empirical and proposed theoretical criteria, the same procedure as the box size experiment was used again, but using the mask diameter and box size decided by the empirical protocol (Table S9). For the mask diameter, the measured diameter of the positive particle density area of the longest particle view was increased by ∼ 10%. Then, the box size was set to approximately twice the mask diameter.

To compare the processing times, all computations of the methemoglobin dataset in the mask diameter and box size experiments were processed with the same single desktop computer (AMD Ryzen Threadripper 2990WX, AMD, Inc.; 32 cores, 3.0 GHz clock time, 132 GB DDR4 memory) equipped with four GPU cards (GeForce RTX 2080 Ti, NVIDIA Corp.).

## Acknowledgements

The authors thank Takahide Yamaguchi and Takamitsu Kohzuma (Ibaraki University) for providing the native nitrite reductase sample; Chiho Masuda and Rieko Sukegawa for assistance with the cryo-EM study; Shinji Aramaki (TVIPS GmbH) for helpful discussion, and Anne Kohtz for English editing and comments. This study was supported by the Platform Project for Supporting Drug Discovery and Life Science Research [Basis for Supporting Innovative Drug Discovery and Life Science Research (BINDS)] from AMED under Grant Number JP20am0101071.

## Author Contributions

T.M. and T.S. conceived and designed the experiments. N.A. and M.K. prepared cryo-EM samples and collected data at the cryo-EM facility in KEK. T.M. processed cryo-EM data. T.M., K.K., A.S., Y.Y., and F.Y. established the environment for the cryo-EM data collection and analysis at the cryo-EM facility in KEK. T.M. and T.S. prepared the manuscript.

## Additional information

Supplementary Information accompanies this paper at http://

### Competing interests

The authors declare no competing financial interests. Reprints and permission information is available online at http://

How to cite this article:

## Materials & Correspondence

All data needed to evaluate the conclusions in the paper are present in the paper and/or the Supplementary Materials. Additional data related to this paper may be requested from the authors.

