## Supplementary Materials for "Size matters: optimal mask diameter and box size for single-particle cryogenic electron microscopy"

Contains:

Supplementary Tables S1-S9

Supplementary Figures S1-S5

**Table S1. Resolutions of the mask diameter (Ø) experiment with nitrite reductase** (pixel size 0.88 Å/pixel, box size 428 pixels = 377 Å)

| *Mask Ø (Å)* | *Initial Res. (Å)* | **Polish03 Res. (Å)* |
|---|---|---|
| (0) 100 | 3.6244 | 3.0991 |
| (1) 132 | 3.3943 | 2.9908 |
| (2) 164 | 3.3413 | 2.8512 |
| (3) 196 | 3.2647 | 2.9293 |
| ***Worst - Best*** | 0.3597 | 0.2479 |

**Table S2. Resolutions of the mask diameter (Ø) experiment with muscle aldolase** (pixel size 0.91 Å/pixel, box size 448 pixels = 408 Å)

| *Mask Ø (Å)* | *Initial Res. (Å)* | *Polish03 Res. (Å)* |
|---|---|---|
| (0) 124 | 2.6821 | 2.5165 |
| (1) 142 | 2.6999 | 2.5165 |
| (2) 160 | 2.6302 | 2.5011 |
| (3) 196 | 2.6133 | 2.5011 |
| (4) 232 | 2.6133 | 2.5011 |
| ***Worst - Best*** | 0.0866 | 0.0154 |

**Table S3. Resolutions of the mask diameter (Ø) experiment with methemoglobin** (pixel size 0.556 Å/pixel, box size 480 pixels = 267 Å)

| *Mask Ø (Å)* | *Initial Res. (Å)* | *Ctf03 Res. (Å)* |
|---|---|---|
| (0) 76 | 3.1398 | 2.8093 |
| (1) 98 | 2.9328 | 2.7513 |
| (2) 120 | 2.9328 | 2.7233 |
| (3) 142 | 2.8697 | 2.7233 |
| ***Worst - Best*** | 0.2701 | 0.0860 |

**Table S4 Processing times relative to mask diameter (Ø) with methemoglobin**
(pixel size 0.556 Å/pixel, box size 480 pixels = 267 Å)

| ***MaskØ (Å)** | ***Initial (hours)*** | ***Ctf03 (hours)*** |
|---|---|---|
| (0) 76 (137) | 0.82 | 1.07 |
| (1) 98 (176) | 0.88 | 1.05 |
| (2) 120 (216) | 0.75 | 1.42 |
| (3) 142 (255) | 0.80 | 1.35 |

*The values in parentheses are in pixels.

**Table S5. Resolutions of the box size experiment with nitrite reductase**
(pixel size 0.88 Å/pixel, mask Ø 114 pixels = 100 Å)

| ***Defocus Percentile*** | ***(a) 100*** | ***(b) ~75*** | ***(c) ~50*** | | | |
|---|---|---|---|---|---|---|
| ***Step ID*** | ***Res. (Å)*** | ***Res. (Å)*** | ***Res. (Å)*** | ***(c) – (a)*** | ***(c) – (b)*** | ***(b) – (a)*** |
| Initial | 3.34125 | 3.28078 | 3.36239 | *0.02114* | *0.08161* | *-0.06047* |
| Ctf01 | 3.21564 | 3.21829 | 3.36239 | *0.14675* | *0.14410* | *0.00265* |
| Polish01 | 2.99077 | 3.01714 | 3.12889 | *0.13812* | *0.11175* | *0.02637* |
| Ctf02 | 2.90939 | 2.96421 | 3.08603 | *0.17664* | *0.12182* | *0.05482* |
| Polish02 | 2.88973 | 2.96421 | 3.08603 | *0.19630* | *0.12182* | *0.07448* |
| Ctf03 | 2.90939 | 2.93843 | 3.08603 | *0.17664* | *0.14760* | *0.02904* |
| Polish03 | 2.87034 | 2.96421 | 3.08603 | *0.21569* | *0.12182* | *0.09387* |
| ***Initial - Best*** | *0.47091* | *0.34235* | *0.27636* | | | |

**Table S6. Resolutions of the box size experiment with muscle aldolase**
(pixel size 0.91 Å/pixel, mask Ø 136 pixels = 124 Å)

| ***Defocus Percentile*** | ***(a) 100*** | ***(b) ~75*** | ***(c) ~50*** | | | |
|---|---|---|---|---|---|---|
| ***Step ID*** | ***Res. (Å)*** | ***Res. (Å)*** | ***Res. (Å)*** | ***(c) – (a)*** | ***(c) – (b)*** | ***(b) – (a)*** |
| Initial | 2.61333 | 2.58844 | 2.70400 | *0.09067* | *0.11556* | *-0.02489* |
| Ctf01 | 2.53217 | 2.56000 | 2.59288 | *0.06071* | *0.03288* | *0.02783* |
| Polish01 | 2.51654 | 2.53217 | 2.55784 | *0.04130* | *0.02567* | *0.01563* |
| Ctf02 | 2.50110 | 2.53217 | 2.55784 | *0.05674* | *0.02567* | *0.03107* |
| Polish02 | 2.50110 | 2.50495 | 2.55784 | *0.05674* | *0.05289* | *0.00385* |
| Ctf03 | 2.50110 | 2.47830 | 2.55784 | *0.05674* | *0.07954* | *-0.02280* |
| Polish03 | 2.50110 | 2.47830 | 2.55784 | *0.05674* | *0.07954* | *-0.02280* |
| ***Initial - Best*** | *0.11223* | *0.11014* | *0.14616* | | | |

**Table S7. Resolutions of the box size experiment with methemoglobin**
(pixel size 0.556 Å/pixel, mask Ø 138 pixels = 76 Å)

| ***Defocus Percentile*** | ***(a) 100*** | ***(b) ~75*** | ***(c) ~50*** | | | |
|---|---|---|---|---|---|---|
| ***Step ID*** | ***Res. (Å)*** | ***Res. (Å)*** | ***Res. (Å)*** | ***(c) – (a)*** | ***(c) – (b)*** | ***(b) – (a)*** |
| Initial | 2.93275 | 2.92107 | 2.96533 | *0.03258* | *0.04426* | *-0.01168* |
| Ctf01 | 2.86968 | 2.75651 | 3.03766 | *0.16798* | *0.28115* | *-0.11317* |
| Polish01 | 2.72327 | 2.75651 | 2.89637 | *0.17310* | *0.13986* | *0.03324* |
| Ctf02 | 2.72327 | 2.75651 | 2.83055 | *0.10728* | *0.07404* | *0.03324* |
| Polish02 | 2.75134 | 2.71822 | 2.83055 | *0.07921* | *0.11233* | *-0.03312* |
| Ctf03 | 2.72327 | 2.75651 | 2.83055 | *0.10728* | *0.07404* | *0.03324* |
| Polish03 | 2.75134 | 2.75651 | 2.83055 | *0.07921* | *0.07404* | *0.00517* |
| ***Initial - Best*** | *0.20948* | *0.20285* | *0.13478* | | | |

**Table S8. Processing times relative to box size with methemoglobin**
(pixel size 0.556 Å/pixel)

| ***Defocus Percentile*** | ****Box Size (Å)*** | ****MaskØ (Å)*** | ***Padding*** | ***Total (hours)*** |
|---|---|---|---|---|
| ***(a) 100*** | 267 (480) | 120 (216) | OFF | 24.47 |
| ***(b) ~75*** | 196 (352) | 120(216) | OFF | 12.66 |
| ***(c) ~50*** | 125 (224) | 120 (216) | OFF | 7.82 |
| ***(c) ~50*** | 125 (224) | 120 (216) | ON | 11.24 |
| ***Empirical*** | 178 (320) | 84 (151) | OFF | 9.50 |

*The values in parentheses are in pixels.

**Table S9. Parameter settings for the direct comparison between empirical and proposed theoretical criteria**

| | |
|---|---|
| ***Nitrite Reductase*** | *Pixel size 0.88 Å/pixel* |
| *Positive Ø* | 114 pixels (100 Å) |
| *Mask Ø* | 125 pixels (110 Å) |
| *Box Size* | 256 pixels (225 Å) |
| *Box Size / Mask Ø* | 2.05 |
| ***Muscle Aldolase*** | *Pixel size 0.91 Å/pixel* |
| *Positive Ø* | 136 pixels (124 Å) |
| *Mask Ø* | 152 pixels (138 Å) |
| *Box Size* | 320 pixels (291 Å) |
| *Box Size / Mask Ø* | 2.11 |
| ***Methemoglobin*** | *Pixel size 0.556 Å/pixel* |
| *Positive Ø* | 137 pixels (76 Å) |
| *Mask Ø* | 151 pixels (84 Å) |
| *Box Size* | 320 pixels (178 Å) |
| *Box Size / Mask Ø* | 2.12 |

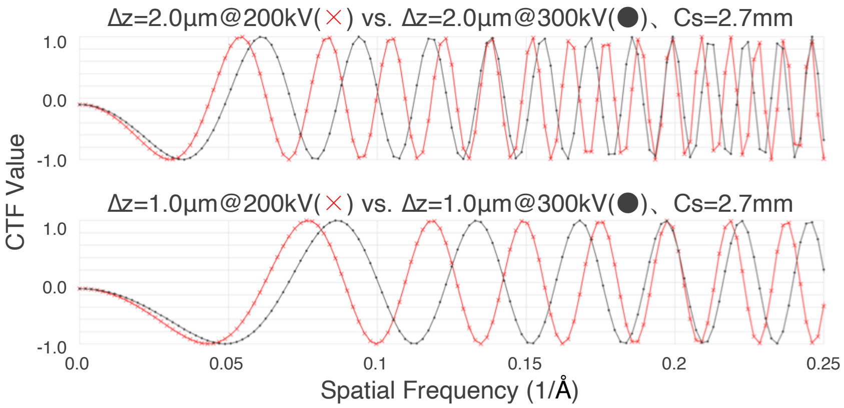


**Fig. S1. CTF differences of the same defocus values between 200 kV and 300 kV**

Examples demonstrating CTF curve differences of the same defocus values between 200 kV and 300 kV acceleration voltage. The 1D CTF curves were simulated with a pixel size of 1.0 Å/pixel and a box size of 512 pixels. (Top) For both 200 kV (red) and 300 kV (black), spherical aberration (Cs) and defocus value were 2.7 mm and 2.0 µm, respectively. (Bottom) Same as A but defocus was 1.0 µm. The oscillations of 200 kV CTF curves are more rapid than of 300 kV curves, especially at the high-frequency range, resembling the difference between CTF curves of large (top) and small (bottom) defocus values, respectively. Also, there are more zero-crossing points where no information is transferred in the CTF curves of a large-defocus example than of a small-defocus example. Approximately 1.27 times the defocus value of 200 kV is the "equivalent defocus" for 300 kV. Therefore, the CTF limit and the PSF width tend to require a larger mask diameter and box size for 200 kV than for 300 kV.

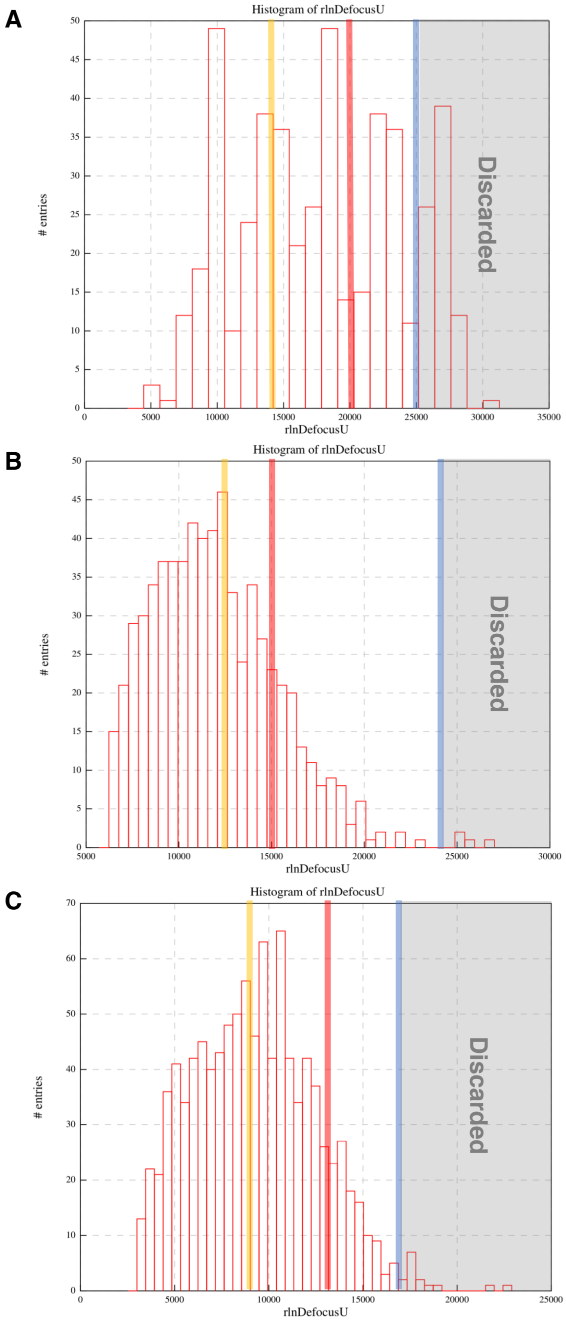


**Fig. S2. Defocus distributions of the datasets**

The defocus distribution histograms of (A) nitrite reductase, (B) muscle aldolase, and (C) methemoglobin datasets. Vertical lines indicate the positions of the maximum (blue), ~75$^{th}$ percentile (red), and ~50$^{th}$ percentile (yellow) defocus values, after discarding the micrographs having too-large defocus values (gray area).

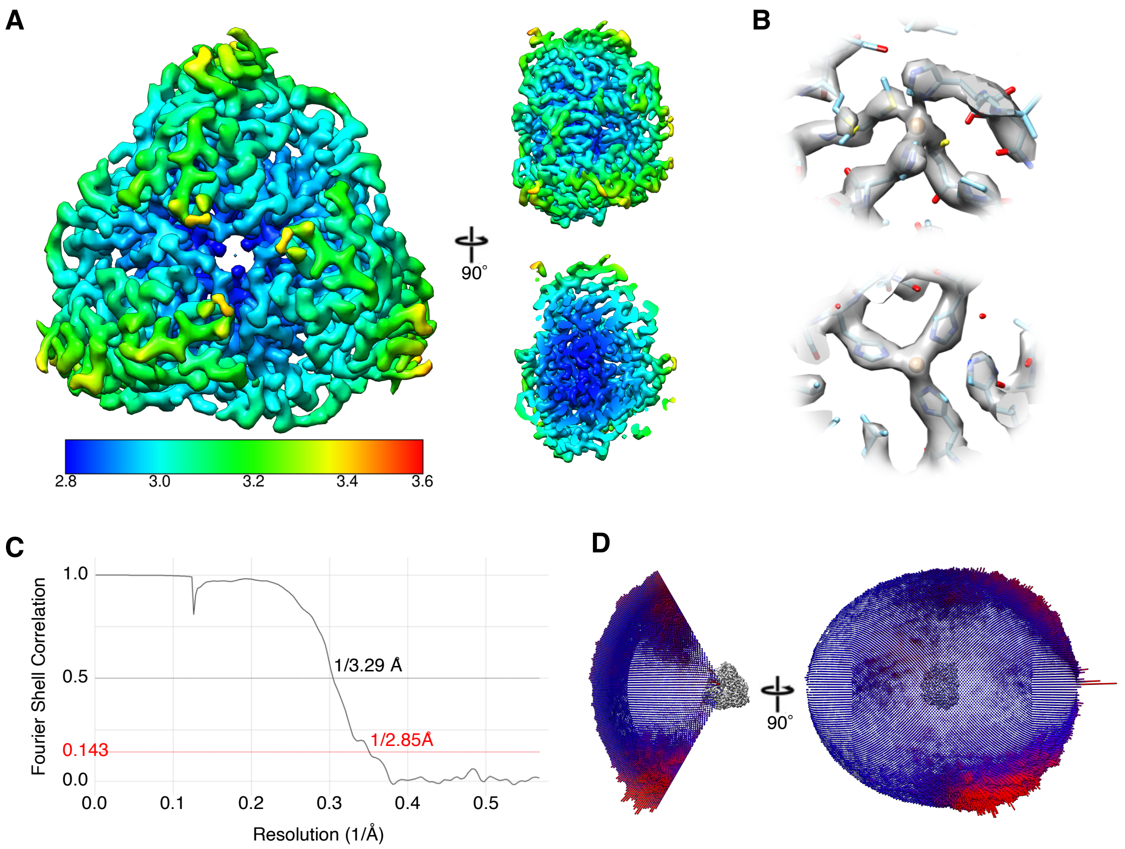


**Fig. S3. The final result of the nitrite reductase dataset**

Validation of the SPA result of the nitrite reductase dataset. (A) Final cryo-EM reconstruction locally filtered and colored by estimated local resolution, ranging from ~2.8 to ~3.6 Å. (B) Zoomed-in view of the 3D density map (gray) at type 1 (top) and type 2 (bottom) Cu ion active sites, with the fitted atomic model (EMD-0731, PDB ID 6KNG). (C) FSC curve showing the resolution of 3.29 Å at 0.5 FSC and 2.85 Å at 1.43 FSC. (D) Orientation distributions.

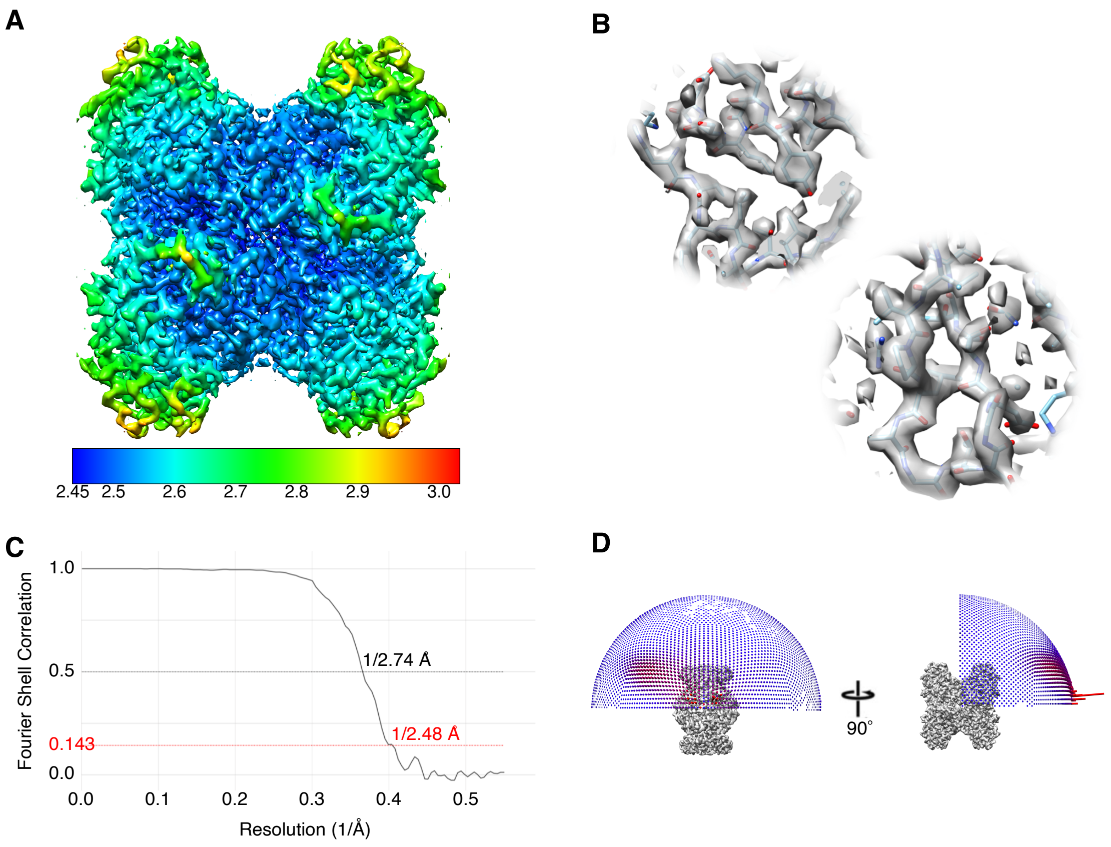


**Fig. S4. The final result of the muscle aldolase dataset**

Validation of the SPA result of the muscle aldolase dataset. (A) Final cryo-EM reconstruction locally filtered and colored by estimated local resolution, ranging from ~2.45 to ~3.0 Å. (B) Zoomed-in view of the 3D density map (gray) with the fitted atomic model (PDB ID 5VY5). (C) FSC curve showing the resolution of 2.74 Å at 0.5 FSC and 2.48 Å at 1.43 FSC. (D) Orientation distributions.

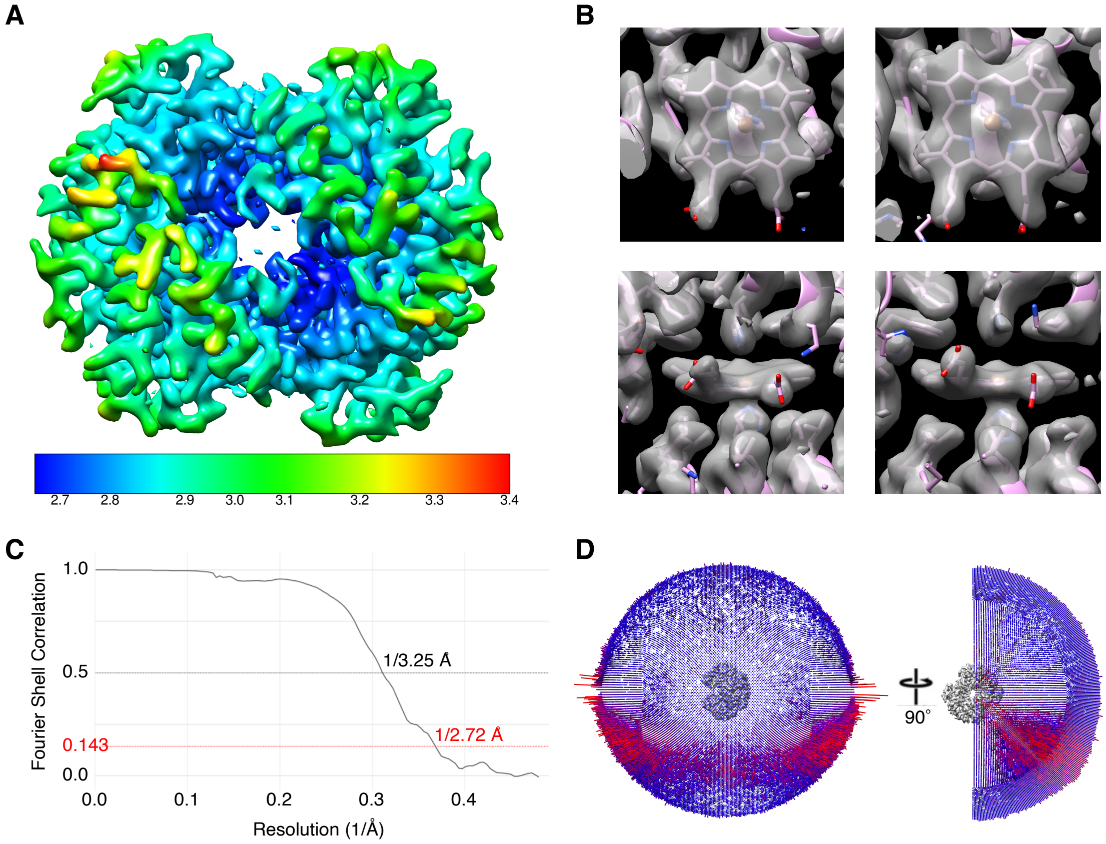


**Fig. S5. The final result of the methemoglobin dataset**

Validation of the SPA result of the methemoglobin dataset. (A) Final cryo-EM reconstruction locally filtered and colored by estimated local resolution ranging from ~2.7 to ~3.4 Å. (B) Zoomed-in view of the 3D density map (gray) of the heme cofactors from subunit α1 (left) and β2 (right) with the fitted atomic model (PDB ID 6NBC). (C) FSC curve showing the resolution of 3.25 Å at 0.5 FSC and 2.72 Å at 1.43 FSC. (D) Orientation distributions.
